# Cytoplasmic TDP-43 accumulation drives changes in C-bouton number and size in a mouse model of sporadic Amyotrophic Lateral Sclerosis

**DOI:** 10.1101/2022.05.20.492885

**Authors:** Anna Normann Bak, Svetlana Djukic, Marion Kadlecova, Thomas Hartig Braunstein, Dennis Bo Jensen, Claire Francesca Meehan

**Affiliations:** Department of Neuroscience, University of Copenhagen; Department of Biomedical Sciences, University of Copenhagen

## Abstract

An altered neuronal excitability of spinal motoneurones has consistently been implicated in Amyotrophic Lateral Sclerosis (ALS) leading to several investigations of synaptic input to these motoneurones. One such input that has repeatedly been shown to be affected is a population of large cholinergic synapses terminating mainly on the soma of the motoneurones referred to as C-boutons. Most research on these synapses during disease progression has used transgenic Superoxide Dismutase 1 (SOD1) mouse models of the disease which have not only produced conflicting findings, but also fail to recapitulate the key pathological feature seen in ALS; cytoplasmic accumulations of TAR DNA-binding protein 43 (TDP-43). Additionally, they fail to distinguish between slow and fast motoneurones, the latter of which have more C-boutons but are lost earlier in the disease.

To circumvent these issues, we quantified the frequency and volume of C-boutons on traced soleus and gastrocnemius motoneurones, representing predominantly slow and fast motor pools respectively. Experiments were performed using the TDP-43ΔNLS mouse model that carries a transgenic construct of TDP-43 devoid of its nuclear localization signal, preventing its nuclear import. This results in the emergence of pathological TDP-43 inclusions in the cytoplasm, modelling the main pathology seen in this disorder, accompanied by a severe and lethal ALS phenotype.

Our results confirmed changes in both the number and volume of C-boutons with a decrease in number on the more vulnerable, predominantly fast gastrocnemius motoneurones and an increase in number on the less vulnerable, predominantly slow soleus motoneurones. Importantly, these changes were only found in male mice. However, both sexes and motor pools showed a decrease in C-bouton volume. Our experiments confirm that cytoplasmic TDP-43 accumulation is sufficient to drive C-bouton changes.

## Introduction

Excitotoxicity has long been proposed as a mechanism contributing to motoneurone death in the neurodegenerative disease Amyotrophic Lateral Sclerosis (ALS) (Choi, 1992; Leigh and Meldrum, 1996). Consequently, much research has focused on changes in both the intrinsic excitability (Meehan *et al*., 2010; Jensen *et al*., 2020a; Jensen *et al*., 2021; Jorgensen *et al*., 2021) and synaptic excitation of the motoneurones (Broadhead *et al*., 2021; Allodi *et al*., 2021; Nagao *et al*., 1998; Ikemoto *et al*., 2002; Zang *et al*., 2005; Schütz, 2005; Sunico *et al*., 2011; Casas *et al*., 2013). One particular excitatory synaptic input that has received considerable attention is the C-bouton (Herron and Miles, 2012; Milan *et al*., 2015; Konsolaki *et al*., 2020; Soulard *et al*., 2020). These large cholinergic synapses are mostly found on the soma and proximal dendrites of spinal motoneurones and function to increase the gain control of motoneurones in a task-dependent manner, particularly in situations of strong physiological drive to the motoneurones (Miles *et al*., 2007; Zagoraiou *et al*., 2009; Witts *et al*., 2014). Consistent with this role, they are found to be more densely distributed on fast fatigable motoneurones that are recruited only when considerable force is required (Hellström *et al*., 2003). Importantly, these are the class of motoneurones that are widely accepted to be the most vulnerable in ALS (Frey *et al*., 2000; Pun *et al*., 2006; Hegedus *et al*., 2007; Hadzipasic *et al*., 2014; Sharma *et al*., 2016; Spiller *et al*., 2016). Alterations in the number of C-boutons on spinal motoneurones have been observed in post mortem tissue from sporadic ALS patients (sALS) (Nagao *et al*., 1998). However, these represent end-stage changes when the large majority of fast motoneurones are lost (Nijssen *et al*., 2017). Consequently, animal models of the disease are the only way to study changes in these synapses during the disease progression, but such studies have yielded inconsistent results (Dukkipati *et al*., 2017).

The most commonly used animal model to study the changes in C-boutons in ALS has been the G93A SOD1 mouse model (Tu *et al*., 1996; Gurney, 1997). This model is based on a hereditary type of ALS where a toxic gain of function in the SOD1 enzyme results in the development of an ALS phenotype (Gurney *et al*., 1994). Although SOD1 mutations account for ∼12-23% of familial ALS (fALS) (Andersen, 2006; Chiò *et al*., 2008), this equates to only 1-2% of all ALS cases, as fALS only accounts for 5-10% of these (Brown and Al-Chalabi, 2017; Mathis *et al*., 2019). Additionally, SOD1 mouse models of the disease, although developing a clear motoneurone disease phenotype, do not recapitulate the major pathological feature of ALS observed in 97% of patients; namely cytoplasmic accumulation of TDP-43 (Neumann *et al*., 2006; Scotter *et al*., 2015; Gao *et al*., 2018; Prasad *et al*., 2019). Despite this, the G93A SOD1 model has almost exclusively been used to investigate C-bouton changes at different stages of disease progression.

Reductions in the number of C-boutons on spinal motoneurones have been reported in SOD1 mice, but this appears to be mostly restricted to the disease end-stage (Gallart-Palau *et al*., 2014; Milan *et al*., 2015; Dukkipati *et al*., 2017; Andres-Benito *et al*., 2019). Although, it should be noted that this time point is normally the ethical humane endpoint, and so not directly comparable with human disease end-stage. Other studies have reported a gain in the number of C-boutons in symptomatic mice (Casas *et al*., 2013; Gallart-Palau *et al*., 2014), while some found no change in the overall number at any disease stage, but instead report a change in the size of the C-boutons (Pullen and Athanasiou, 2009; Herron and Miles, 2012; Aliaga *et al*., 2013; Saxena *et al*., 2013). Changes observed with respect to size have also not been consistent. Most studies have tended to show an increase in C-bouton size that increases further with disease progression (Herron and Miles, 2012; Aliaga *et al*., 2013; Saxena *et al*., 2013). However, other studies have reported no change or even a decrease in size at different disease stages (Pun *et al*., 2006; Pullen and Athanasiou, 2009; Milan *et al*., 2015).

These inconsistencies have been proposed to be due to differences in the way data was collected and analysed statistically (Dukkipati *et al*., 2017), but may also reflect the broader study design including disease stage and sex of the mice. Despite an early study reporting an increase in C-bouton size only in male SOD1 mice (Herron and Miles, 2012), very few subsequent studies have systematically investigated or controlled for this sex difference. Additionally, none have controlled for a loss of fast motoneurones with disease progression. Most studies used immunohistochemical labelling of choline acetyltransferase to identify spinal motoneurones, which does not allow a distinction between slow and fast motoneurones. Therefore, reports of a reduction in the number of C-boutons may simply reflect the earlier loss of the fast motoneurones, which normally possess more C-boutons (Conradi *et al*., 1979a; Kellerth *et al*., 1979; Conradi *et al*., 1979b; Hellström *et al*., 2003). This also fails to distinguish between motoneurones that are still functionally connected at the neuromuscular junction as the disease progresses and those that are not. This is a crucial distinction to make as a loss of function at the neuromuscular junction has been shown to cause a decrease in the size of C-boutons (Casanovas *et al*., 2017; Jensen *et al*., 2020b), whilst axotomy has been demonstrated to induce even more severe changes with a drastic reduction in both the number and size of C-boutons (Salvany *et al*., 2019).

To address these issues, we performed experiments in a relatively new inducible mouse model of sporadic ALS (TDP43ΔNLS) which recapitulates the key pathological feature observed in the disease that is cytoplasmic accumulations of TDP-43 (Neumann *et al*., 2006; Scotter *et al*., 2015; Gao *et al*., 2018; Prasad *et al*., 2019). This mouse contains a TDP gene with a defective nuclear localisation signal, the expression of which is controllable using the tetracycline transactivator system. To specifically control for the factors mentioned above, retrograde tracing was employed with different tracers injected into the predominantly slow soleus muscle and the predominantly fast gastrocnemius muscle in both male and female mice. This allows, not only a distinction between predominantly fast and slow motoneurones, but also requires functional connectivity at the neuromuscular junction as the tracer has to be transported retrogradely from the muscle to the soma.

## Methods

The experimental procedures were approved by the Danish Animal Experiments Inspectorate (Permission number 2018-15-0201-01426) and were in accordance with the EU Directive 2010/63/EU.

### Mice

For these experiments, TDP43ΔNLS mice were obtained by crossing tetO-hTDP-43-ΔNLS line 4 (https://www.jax.org/strain/014650) with NEFH-tTA line 8 (https://www.jax.org/strain/025397). Bigenic mice carry a tetracycline-repressible TARDBP gene construct that lacks the nuclear localization signal, which is located downstream of a tetracycline response element composed of TetO operator sequences. Hence, when the animals are kept on a diet containing the tetracycline-analogue Doxycycline, this chemical compound binds to a constitutively expressed tetracycline-transactivator (tTA) protein, blocking the interaction between tTA and the above-mentioned TetO promoter - and thereby represses the transcriptional activation of the ALS-causing TDP-43 ΔNLS. However, when the animals are switched from Dox-containing food to normal food, the expression of this toxic, mislocalized protein is turned on as tTA is free to drive promoter activation. This results in a mouse line with a regulatable expression of cytoplasmic-insoluble human TDP-43 in large-caliber axons of the brain and spinal cord including motoneurones.

Genotyping was performed by extracting DNA from ear clippings from all mice, which were then genotyped using PCR and four sets of transgene specific primers. Two sets of primers, one for Tg(tetO-TARDBP), the human TDP-43, and one for Tg(NEFH-tTA)8Vle, the promoter, and two sets of internal control primers, one for each transgene. Primer sequence used for the Tg(tetO-TARDBP) were forward 5’- TTG CGT GAC TCT TTA GTA TTG GTT TGA TGA-3’ and reverse 5’- CTC ATC CAT TGC TGC TGC G-3’ and for the Tg(NEFH-tTA)8Vle forward 5’- CTC GCG CAC CTG CTG AAT-3’ and reverse 5’- CAG TAC AGG GTA GGC TGC TC-3’. The control primers were respectively 5’- CAA ATG TTG CTT GTC TGG TG-3’ with 5’- GTC AGT CGA GTG CAC AGT TT-3’ and 5’- CTA GGC CAC AGA ATT GAA AGA TCT-3’ with 5’-GTA GGT GGA AAT TCT AGC ATC ATC C-3’. Primers were as recommended by The Jackson Laboratory. Only mice expressing both transgenes were used for experiments.

### Behavioural characterisation

The animals were housed in our own local animal rooms at the Panum Institute serviced by the Department of Experimental Medicine at the Faculty of Health at Copenhagen University. Bigenic mice were maintained on doxycycline in chow (200mg/kg SAFEDIET) until 7 weeks of age at which point the doxycycline food was replaced with standard chow. To characterise the disease progression after transgene induction in our colony and to identify a good time point to perform experiments, an initial cohort of six mice were used. In these mice, a simple grip test was performed in which the mice were placed on a cage lid which was then inverted over a soft surface, and the endurance time that the mice could hold on was measured for each mouse. From this, a time point was selected for tracing experiments in a second cohort of induced mice. Bigenic mice of the same age were maintained on doxycycline as controls. 10 induced mice (5 males, 5 females) and 10 aged matched non-induced mice (5 males, 5 females) were used for the main experiments.

### Tracing of spinal motoneurones

At approximately 10 1/2 weeks of age, tracers were injected into the predominantly fast gastrocnemius muscle and the predominantly slow soleus muscle. Prior to surgery mice were administered Temgesic (0.05-0.1mg/kg) for analgesia and the antibiotic Tribrissen (0.1mL of 1:10 dilution from 200/40mg/mL). Under isofluorane anaesthesia, a small incision was made bilaterally on the hindlimb to expose the Achilles tendon and from here gastrocnemius and soleus muscles were identified. The tracer Cholera toxin subunit B conjugated to Alexa Fluor 488, (CTB 488, Thermo Fisher Scientific 0.05% in 5 μl Phosphate Buffered Saline, PBS) was injected into the gastrocnemius muscles and the tracer Fast Blue tracer (Polysciences cat.no. 17740-5 at 1.5% in 3μL PBS) was injected into the soleus muscles. Post-operative pain control was maintained with Temgesic delivered in a mix with Nutella for the following 2-3 days.

### Immunohistochemistry

At 11 weeks of age (approximately 3-4 days after the tracer injections) the animals were euthanized with Sodium Pentobarbital (120mg/kg), and transcardially perfused with 0.9 % saline followed by 5-6 minutes perfusion with freshly made 4 % paraformaldehyde. The spinal cords were removed and post-fixed for 2 hours in 15 % sucrose in the same fixative and then immersed in 30 % sucrose in PBS overnight for cryoprotection. The lumbar enlargement of the spinal cord was then cut into 50 μm thick horizontal sections and ventral sections were processed for immunohistochemistry. Sections were blocked for 4 hours in 5 % donkey serum in PBST (0.3 %) before incubation in the primary antibody diluted in blocking solution overnight (goat anti-VAChT, Millipore 1:2000 LOT: 2899777 and 3593751 at 1:1000). The tissue was then incubated for 2 hours in the secondary antibody (donkey anti-goat, Alexa Fluor 650 1:1000 Thermo Fisher Scientific Lot: SA2325974), rinsed and then mounted onto glass slides and cover-slipped with ProLong™ Gold Antifade Mountant medium (ThermoFisher Catalog number: P10144).

### Imaging and Analysis

Traced motoneurones were imaged with a Zeiss LSM700 confocal microscope, using a 20x/0.8 Plan Apochromat Air objective and Zeiss ZEN 2.3 Black Software. Analyses were then performed using Z-stacks in both Zeiss Zen 3.2 Blue Software and FiJi (ImageJ, NIH). Blinding was attempted but made difficult by the presence of unusual swellings of the axon hillock and accumulation of tracers in this region in the induced mice. Therefore, an unbiased automated process to measure the C-boutons was developed. In order to do full cell analyses only cells with somas that were completely contained within the tissue section were used. The cells and their respective C-boutons were segmented using ZEN Software where a neural network (Zeiss Intellesis) was trained to recognize traced green cells, traced blue cells and C-boutons in their respective channels. From these binary images were created consisting of only cell and background or C-boutons and background. The binary images were analysed in FiJi using a macro that made the stack dimensions isotropic and then used a function to grow the soma of the cell in 3D, to include the surrounding C-boutons, and to ensure that only these would be included in the analysis. The FiJi “3D Objects Counter” (Bolte and Cordelières, 2006) was used to analyse the cell and the boutons respectively (see the process illustrated in figure 1).

**Figure 1:**
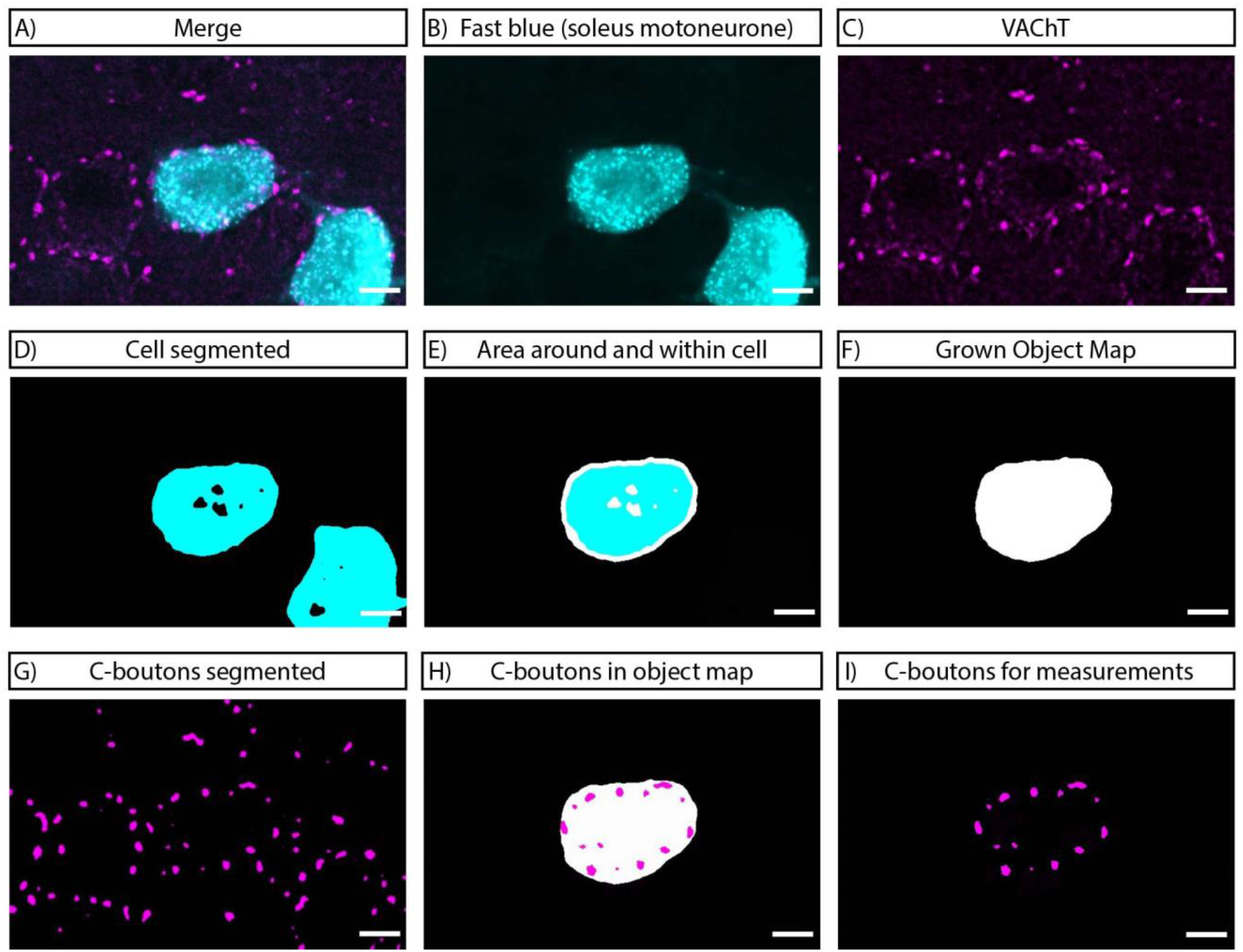
Example of the automated analysis of C-boutons. A) Merged illustration showing a B) Fast Blue traced soleus motoneurone and C) C-boutons stained with antibodies against VAChT. D) Cell soma segmented by a trained neuronal network (Intellesis trainable segmentation) to a binary picture consisting only of cell soma and background. E) Area around and within cell included by running a “distance transform” function in Image J creating an object map seen in F). G) C-boutons segmented by a trained neuronal network (Intellesis trainable segmentation) to a binary picture consisting only of C-boutons and background. H) Object map from the cell soma applied onto the segmented C-boutons to isolate the ones needed for analysis. I) The final C-boutons for analysis by 3D Object Counter. Scale bar is 10 μm.

Images for the figures were prepared using Fiji (Image J, NIH) and Adobe Illustrator. Brightness, contrast, gaussian blur and background subtraction were adjusted for image presentation using Image J and this was performed uniformly across the entire image.

### Statistical Analysis

The data were analysed by cell, with each cell constituting a separate “n” (as recommended by (Dukkipati *et al*., 2017)) as this takes into consideration the heterogeneous nature of the motoneurone population within a single animal. This also allowed us to observe if the overall ranges were altered and to control for soma size. GraphPad Prism software was used for all statistical analysis. Normality tests (D’Agostino & Pearson omnibus normality test) were used to determine whether parametric statistics or non-parametric tests should be used. Statistical significance was accepted at the P≤0.05 level. In the figures, stars are used to indicate significant differences as follows: * (P ≤0.05), ** (P ≤ 0.01), ***(P≤0.001) and ****(P≤0.0001). Full details of the statistical tests are provided in Table 1 and 2.

**Table 1:** Statistical analysis of the data presented in figure 3.

| Parameter | MN Pool | Sex | Induction Status | Mean (SD) or median (IQR) | N cells (n for mice) | Statistic test | Test value | P value |
| --- | --- | --- | --- | --- | --- | --- | --- | --- |
| Number of C-boutons /Cell | Soleus | Both | Non Induced | 51.27 (13.33) | 66 (10) | T test | t=5.562 | <0.0001 |
|  | Gastrocnemius | Both | Non induced | 64.43 (14.67) | 76 (9) |  |  |  |
| Number of C-boutons/Cell | Soleus | Female | Non Induced | 53.0 (13.44) | 33 (5) | T test | t=2.071 | 0.0421 |
|  | Gastrocnemius | Female | Non Induced | 59.51 (13.18) | 39 (5) |  |  |  |
| Number of C-boutons/Cell | Soleus | Male | Non Induced | 49.55 (13.19) | 33 (5) | T test | t=6.024 | <0.0001 |
|  | Gastrocnemius | Male | Non Induced | 69.62 (14.53) | 37 (4) |  |  |  |
| C-bouton density/Cell (boutons pr. 1000 $\mu\text{m}^2$) | Soleus | Both | Non Induced | 1.982 (0.676) | 66 (10) | Mann-Whitney | U=1494 | <0.0001 |
|  | Gastrocnemius | Both | Non induced | 1.670 (0.603) | 76 (9) |  |  |  |
| C-bouton density/Cell (boutons pr. 1000 $\mu\text{m}^2$) | Soleus | Female | Non Induced | 1.950 (0.683) | 33 (5) | Mann-Whitney | U=314 | 0.0001 |
|  | Gastrocnemius | Female | Non Induced | 1.363 (0.634) | 39 (5) |  |  |  |
| C-bouton density/Cell (boutons pr. 1000 $\mu\text{m}^2$) | Soleus | Male | Non Induced | 2.095 (0.4727) | 33 (5) | Welch's t test | t=2.471 | 0.0166 |
|  | Gastrocnemius | Male | Non Induced | 1853 (0.3267) | 37 (4) |  |  |  |
| Mean C-bouton Volume/Cell ($\mu\text{m}^3$) | Soleus | Both | Non Induced | 83.20 (18.95) | 66 (10) | Welch's t test | t=2.607 | 0.0103 |
|  | Gastrocnemius | Both | Non induced | 75.79 (14.17) | 76 (9) |  |  |  |
| Mean C-bouton Volume/Cell ($\mu\text{m}^3$) | Soleus | Female | Non Induced | 76.27 (17.45) | 33 (5) | T test | t=0.3050 | 0.7612 |
|  | Gastrocnemius | Female | Non Induced | 75.10 (15.00) | 39 (5) |  |  |  |
| Mean C-bouton Volume/Cell ($\mu\text{m}^3$) | Soleus | Male | Non Induced | 90.13 (18.06) | 33 (5) | T test | t=3.609 | 0.0006 |
|  | Gastrocnemius | Male | Non Induced | 76.51 (13.40) | 37 (4) |  |  |  |

**Table 2:** Statistical analysis of the data presented in figure 4 and 5

| Parameter | MN Pool | Sex | Induction Status | Mean (SD) or median (IQR) | N cells (n for mice) | Statistic test | Test value | P value |
| --- | --- | --- | --- | --- | --- | --- | --- | --- |
| Cell Size | Soleus | Both | Non Induced | 25315 (17538) | 66 (10) | Mann-Whitney | U=2263 | 0.7328 |
|  |  |  | Induced | 26633 (12925) | 71 (10) |  |  |  |
|  | Gastrocnemius | Both | Non Induced | 36177 (14164) | 76 (9) | Mann-Whitney | U=2355 | 0.0001 |
|  |  |  | Induced | 31305 (10885) | 98 (10) |  |  |  |
| Cell Size | Soleus | Female | Non Induced | 30900 (16826) | 33 (5) | Welch's t test | t=1.285 | 0.2055 |
|  |  |  | Induced | 26762 (8039) | 36 (5) |  |  |  |
|  | Soleus | Male | Non Induced | 21789 (12518) | 33 (4) | Mann-Whitney | U=489 | 0.2822 |
|  |  |  | Induced | 24593 (13327) | 35 (5) |  |  |  |
| Cell Size | Gastrocnemius | Female | Non Induced | 36222 (17255) | 39 (5) | Mann-Whitney | U=636 | 0.0101 |
|  |  |  | Induced | 31270 (13057) | 48 (5) |  |  |  |
|  | Gastrocnemius | Male | Non Induced | 36802 (9664) | 37 (4) | T test | t=3.073 | 0.0028 |
|  |  |  | Induced | 30813 (8455) | 50 (5) |  |  |  |
| Number of C-boutons/Cell | Soleus | Both | Non Induced | 51.27 (13.33) | 66 (10) | Welch's T test | t=2.086 | 0.0390 |
|  |  |  | Induced | 56.70 (17.04) | 71 (10) |  |  |  |
|  | Gastrocnemius | Both | Non Induced | 64.43 (14.67) | 76 (9) | T test | t=0.7346 | 0.4636 |
|  |  |  | Induced | 62.82 (14.20) | 98 (10) |  |  |  |
| Number of C-boutons/Cell | Soleus | Female | Non Induced | 53.00 (13.44) | 33 (5) | T test | t=0.3217 | 0.7487 |
|  |  |  | Induced | 54.22 (17.63) | 36 (5) |  |  |  |
|  | Soleus | Male | Non Induced | 49.55 (13.19) | 33 (5) | T test | t=2.693 | 0.0090 |
|  |  |  | Induced | 59.26 (16.28) | 35 (5) |  |  |  |
| Number of C-boutons/Cell | Gastrocnemius | Female | Non Induced | 59.51 (13.18) | 39 (5) | T test | t=1.509 | 0.1350 |
|  |  |  | Induced | 63.96 (14.04) | 48 (5) |  |  |  |
|  | Gastrocnemius | Male | Non Induced | 69.62 (14.53) | 37 (4) | T test | t=2.519 | 0.0136 |
|  |  |  | Induced | 61.72 (14.41) | 50 (5) |  |  |  |
| C-bouton density/Cell (boutons pr. 1000 $\mu\text{m}^2$) | Soleus | Both | Non Induced | 1.982 (0.676) | 66 (10) | Mann-Whitney | U=1914 | 0.0648 |
|  |  |  | Induced | 2.025 (0.4368) | 71 (10) |  |  |  |
|  | Gastrocnemius | Both | Non Induced | 1.662 (0.4316) | 76 (9) | T test | t=5.472 | <0.0001 |
|  |  |  | Induced | 2.035 (0.4301) | 98 (10) |  |  |  |
| C-bouton density/Cell (boutons pr. 1000 $\mu\text{m}^2$) | Soleus | Female | Non Induced | 1.950 (0.683) | 33 (5) | Mann-Whitney | U=516 | 0.3541 |
|  |  |  | Induced | 2.046 (0.623) | 36 (5) |  |  |  |
|  | Soleus | Male | Non Induced | 2.095 (0.4727) | 33 (5) | T test | t=1.978 | 0.0521 |
|  |  |  | Induced | 2.347 (0.5698) | 35 (5) |  |  |  |
| C-bouton density/Cell (boutons pr. 1000 $\mu\text{m}^2$) | Gastrocnemius | Female | Non Induced | 1.363 (0.634) | 39 (5) | Mann-Whitney | U=356 | <0.0001 |
|  |  |  | Induced | 2.029 (0.537) | 48 (5) |  |  |  |
|  | Gastrocnemius | Male | Non Induced | 1.853 (0.3267) | 37 (4) | Welch's t test | t=2.333 | 0.0220 |
|  |  |  | Induced | 2.047 (0.4522) | 50 (5) |  |  |  |
| Mean C-bouton Volume/Cell ($\mu\text{m}^3$) | Soleus | Both | Non Induced | 83.75 (27.96) | 66 (10) | Mann-Whitney | U=955 | <0.0001 |
|  |  |  | Induced | 61.56 (20.16) | 71 (10) |  |  |  |
|  | Gastrocnemius | Both | Non Induced | 75.06 (18.40) | 76 (9) | Mann-Whitney | U=2066 | <0.0001 |
|  |  |  | Induced | 63.44 (16.67) | 98 (10) |  |  |  |
| Mean C-bouton Volume/Cell ($\mu\text{m}^3$) | Soleus | Female | Non Induced | 76.27 (17.45) | 33 (5) | T test | t=3.898 | 0.0002 |
|  |  |  | Induced | 61.77 (13.31) | 36 (5) |  |  |  |
|  | Soleus | Male | Non Induced | 92.97 (19.48) | 33 (5) | Mann-Whitney | U=190 | <0.0001 |
|  |  |  | Induced | 61.56 (23.57) | 35 (5) |  |  |  |
| Mean C-bouton Volume/Cell ($\mu\text{m}^3$) | Gastrocnemius | Female | Non Induced | 72.86 (22.10) | 39 (5) | Mann-Whitney | U=551 | 0.0009 |
|  |  |  | Induced | 62.62 (10.70) | 48 (5) |  |  |  |
|  | Gastrocnemius | Male | Non Induced | 582.4 (111.8) | 37 (4) | T test | t=3.776 | 0.0003 |
|  |  |  | Induced | 496.3 (158.5) | 50 (5) |  |  |  |

**Table 3:** Statistical analysis of the regression data presented in figure 4

| Analysis | Induction status | N cells (n for mice) | Test | R<sup>2</sup> | P value | Are the slopes different? | Are the elevations of the slopes different? |
| --- | --- | --- | --- | --- | --- | --- | --- |
| Mean C-bouton Density/ 1000 $\mu\text{m}^2$ for Gastrocnemius motoneurons) | Non Induced | 76 (9) | Linear Regression | 0.2383 | <0.0001 | P=0.9667 | P<0.0001 |
|  | Induced | 98 (10) |  | 0.1008 | <0.0001 |  |  |

## Results

Behavioural tests were used to characterise the disease progression in our colony using an initial cohort of 6 mice. This was used to determine an optimal time to perform experiments. Within 2 weeks of transgene expression, all of the induced mice started to show a motor phenotype. When suspended by their tail, control mice will normally stretch out their hind legs and reach the forelimbs forward to the ground. By 2 weeks post induction, all mice showed a clear clasping phenotype when suspended by their tail. By 4 weeks, all induced mice showed more drastic clasping behaviour (figure 2A).

**Figure 2:**
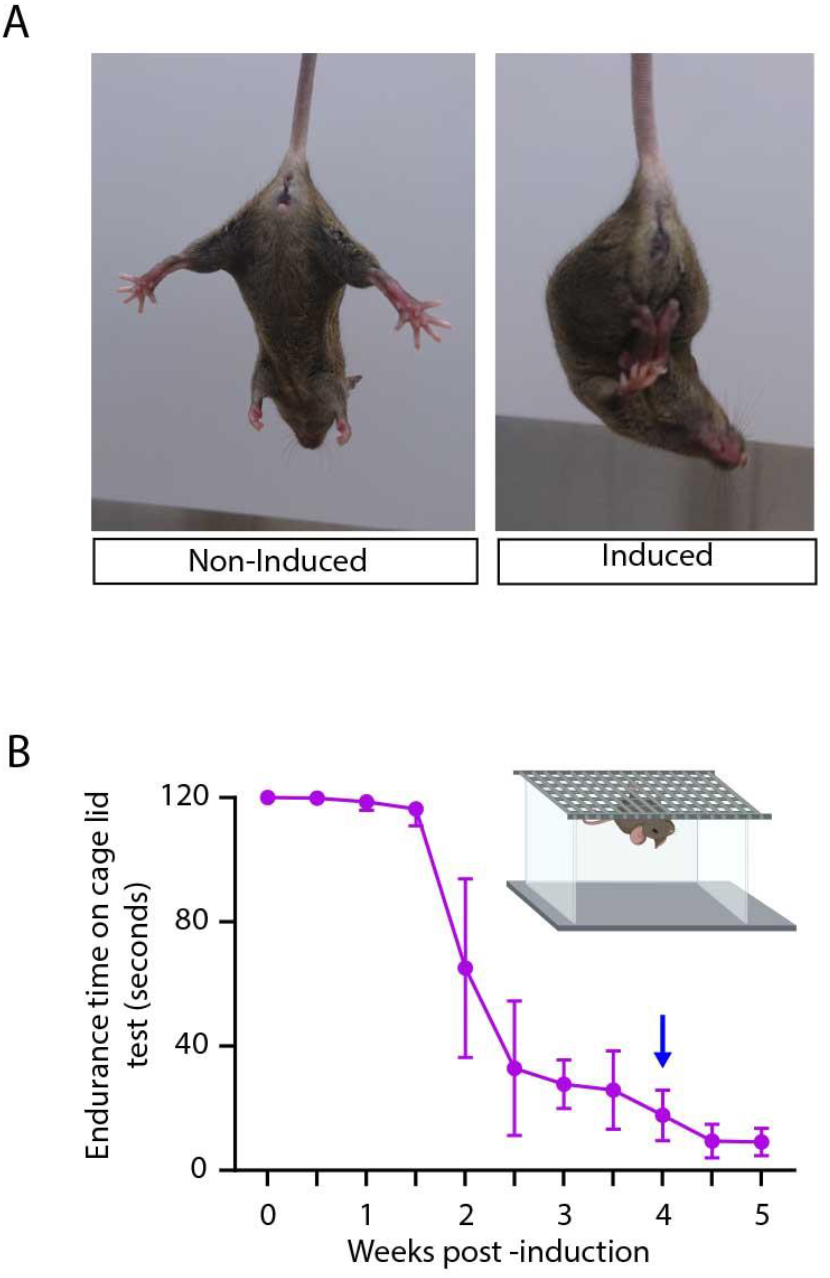
Behavioural phenotype of the animals. **A)** Two age-matched TDP-43 mice carrying both transgenes showing the phenotypical differences between a non-induced control mouse and a mouse at four weeks post induction. **B)** A graph showing the declining grip endurance after induction (blue arrow indicated the time point selected for anatomical experiments). A small illustration shows the principle of the Cage Grip Test.

From 2 weeks post-induction, performance on the cage lid grip test started to deteriorate with mice showing severe impairments by 4 weeks post induction (figure 2B) accompanied by clear muscle atrophy. The mice reached humane endpoint around 5 to 6 weeks post-induction. To investigate the C-boutons at a point prior to end-stage (and for ethical reasons related to recovery surgeries), the anatomical experiments were performed in a second cohort of 10 mice at 4 weeks post-induction.

### Fast motoneurones have a greater number of C-boutons in control animals in both males and females

CTB-488 tracer (green) was used to label gastrocnemius motoneurones and fast blue tracer (blue) was used to label soleus motoneurones in the mice (figure 3A). Examples of C-bouton labelling on both gastrocnemius and soleus motoneurones in the non-induced control group are shown in figure 3B. The mean number of C-boutons per cell was quantified and compared between the predominantly slow soleus- and predominantly fast gastrocnemius motoneurones. Consistent with the previous literature, in the control (non-induced) group, gastrocnemius motoneurones (predominantly fast motoneurones) had significantly more C-boutons per cell than soleus motoneurones (predominantly slow motoneurones) (Conradi *et al*., 1979a; Kellerth *et al*., 1979; Conradi *et al*., 1979b; Hellström *et al*., 2003), an observation that was consistent across both sexes, but was more pronounced in males (figure 3C, table 1). To control for soma size, the average density was calculated (boutons per 1000 μm^2^ surface area), which was significantly different between the two groups (figure 3D, table 1) showing that despite having more C-boutons, the overall density was less in the gastrocnemius motoneurones compared to the soleus motoneurones. This difference was again significant for both sexes (figure 3D, table 1). The mean volume of the C-boutons (mean per cell) was also calculated and this revealed that the volume of C-boutons was significantly larger on soleus motoneurones compared to those on the gastrocnemius motoneurones - but surprisingly, this difference was only significant in males (figure 3E, table 1).

**Figure 3:**
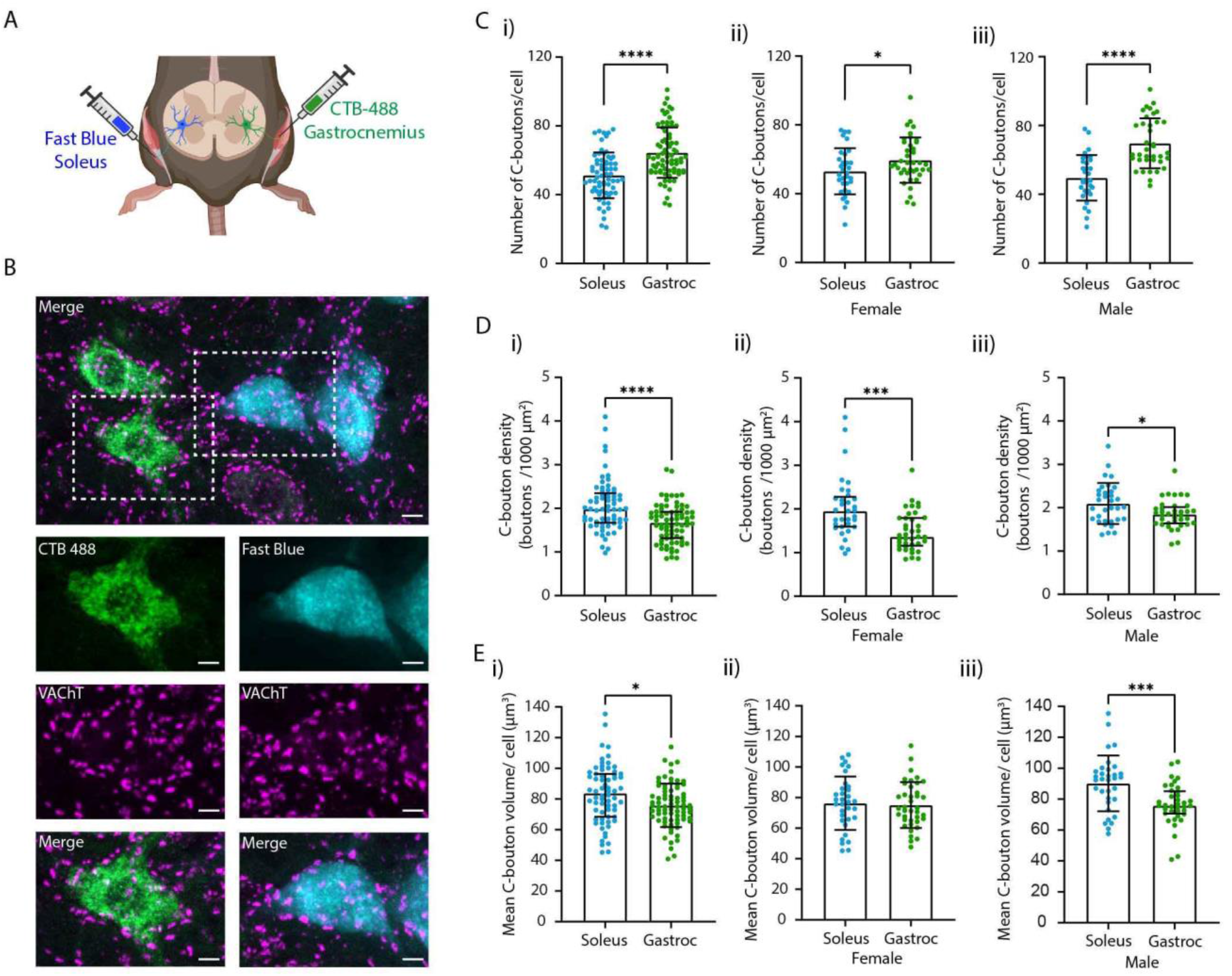
C-bouton parameters on fast and slow motoneurones. **A)** Illustration of the tracer injection into soleus (blue tracer, fast blue) and gastrocnemius (green tracer, CTB conjugated to Alexa 488). **B)** Examples of maximum projections of confocal microscope images of the traced cells that were immunohistochemically stained for VAChT (C-boutons) showing both soleus motoneurones (blue) and gastrocnemius motoneurones (green). Scale bar is 10 μm. **C)** Scatter dot plots showing the number of C-boutons per cell compared between soleus and gastrocnemius motoneurones in control animals with both sexes collapsed (i) then for female (ii) and male (iii) mice separately. **D)** Scatter dot plots showing C-bouton density per cell compared between soleus and gastrocnemius in control animals, with both sexes collapsed (i) then for female (ii) and male (iii) mice separately. **E)** Scatter dot plots showing the average C-bouton volume per cell compared between soleus and gastrocnemius in control animals, with both sexes collapsed (i) then for female (ii) and male (iii) mice separately. In all scatter dot plots the bars represent means with standard deviation (SD) for normally distributed data and medians with interquartile range for non-parametric data.

To control for these observations in induced mice, gastrocnemius- and soleus motoneurones were analysed separately to determine if any differences in the induced animals were due to a loss of one type of motoneurone, or if they were bona fide changes that may be consistent across motoneurone types. Furthermore, any differences were tested to determine if they were consistent across the two sexes.

### TDP-43 mislocation drives an increase in the number of C-boutons in male mice

Examples of C-bouton labelling on both gastrocnemius- and soleus motoneurones from a control and an induced mouse are shown in figure 4A. From these pictures, a slight shrinkage of the gastrocnemius motoneurones can be observed along with an abnormal accumulation of the tracer in the axon hillock. While blinding was attempted, in reality this feature made true blinding difficult and so automated C-bouton identification, quantification and measuring were performed.

**Figure 4:**
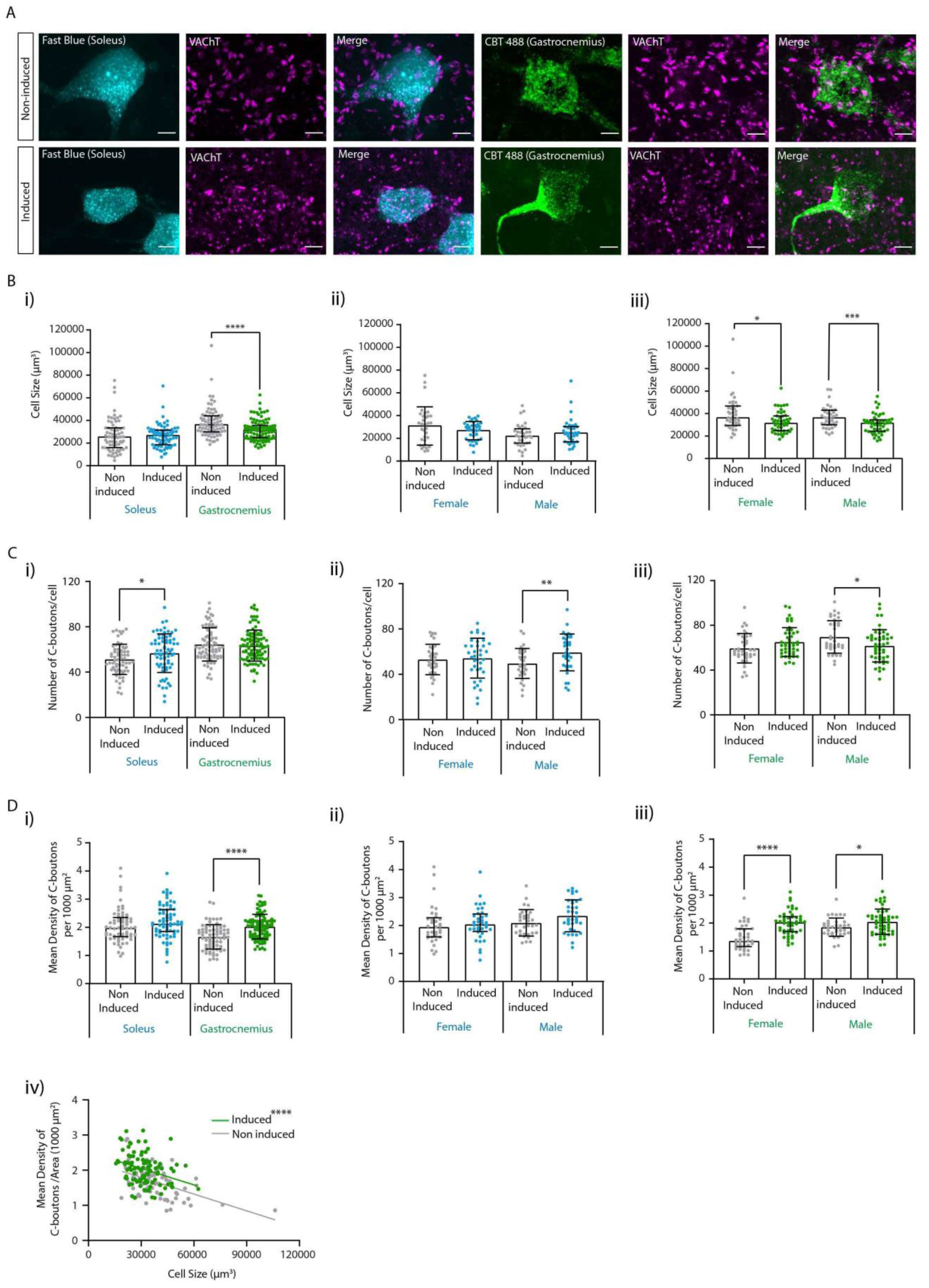
Cytoplasmic TDP-43 drives changes in the number of C-boutons. **A)** Examples of traced motoneurones (soleus: blue, gastrocnemius: green) and immunohistochemically labelled C-boutons (magenta) for a non-induced and an induced male mouse (Scale bar is 10 μm). **B)** Scatter dots plots showing a significant reduction in soma size for gastrocnemius motoneurones only (i). ii and iii show the same data separated into the separate sexes showing that this difference is consistent across both sexes. **C)** Scatter dot plots showing the number of C-boutons per cell. When analysed together, a significant increase in the number of C-boutons is seen for soleus motoneurones only (i). When the data for the different sexes is separated (ii and iii) it can be seen that this difference is only observed for males in addition to a significant decrease in C-bouton number on gastrocnemius motoneurones in males with no changes seen for females. **D)** Scatter dot plots showing the density of C-boutons per cell which is, in fact, significantly higher for gastrocnemius motoneurones due to their reduction in soma size (i).When the same data is analysed separately by sex, there is still no difference for soleus motoneurones (ii) but the significant increase in C-bouton density is consistent across sexes (iii). When the mean density of C-boutons per cell is plotted against soma size (iv), the regression line is significantly higher for induced mice for gastrocnemius motoneurones, showing that the increase in density cannot merely be explained by the reduction in soma size. In all scatter dot plots the bars represent means with standard deviation (SD) for normally distributed data and medians with interquartile range for non-parametric data.

Measurements of the soma size confirmed a significant reduction in cell size for gastrocnemius, but not soleus motoneurones in induced mice compared with non-induced mice for both sexes (Figure 4B). When the total number of C-boutons per cell was compared between non-induced and induced mice, a significant increase in number was observed in induced mice (Figure 4C). This increase appeared to be restricted to the soleus motoneurones. However, further analysis separating data from males and females revealed a significant increase in the mean number of C-boutons per cell for soleus motoneurones, and a decrease in the number for the gastrocnemius motoneurones. However, these changes were only significant in the males with none of these differences observed for females (Figure 4C ii and iii). These changes in the number of C-boutons in induced male mice are unlikely simply to be a consequence of a loss of fast motoneurones as F-tests confirmed no significant differences between the variances of the two groups (non-induced and induced). Furthermore, given our observations in figure 3, a loss of fast motoneurones would be predicted to result in a decrease in the mean number of C-boutons per cell (as these cells normally possess more C-boutons) rather than the increase that we observed on soleus motoneurones.

While a decrease in C-bouton number was observed for gastrocnemius motoneurones, it must also be taken into consideration that these motoneurones were smaller in the induced mice, which could actually contribute to an increased density of C-boutons on these cells. To take this into account, we expressed this as the density of C-boutons per 1000 μm^2^ (figure 4D). This confirmed that the cell shrinkage contributed to a significant increase in the density of C-boutons on gastrocnemius motoneurones on both female and male induced mice. However, this increased density was not entirely proportional to the degree of soma shrinkage, as linear regression revealed that even though C-bouton density was significantly correlated with soma size for both groups, the elevation of the regression line was significantly higher for the induced mice (figure 4D) demonstrating an actual increase in density in the induced mice compared to the controls.

### TDP-43 mislocation drives a decrease in C-bouton volume on both fast and slow motoneurones

The mean C-bouton volume was calculated for each cell. In induced mice, C-boutons were significantly smaller than on the non-induced controls for both soleus and gastrocnemius motoneurones by 26.5 % and 13.6 %, respectively (figure 5A, table 2). This difference was significant for both cell pools in both males and females (figure 5B, C, table 2).

**Figure 5.**
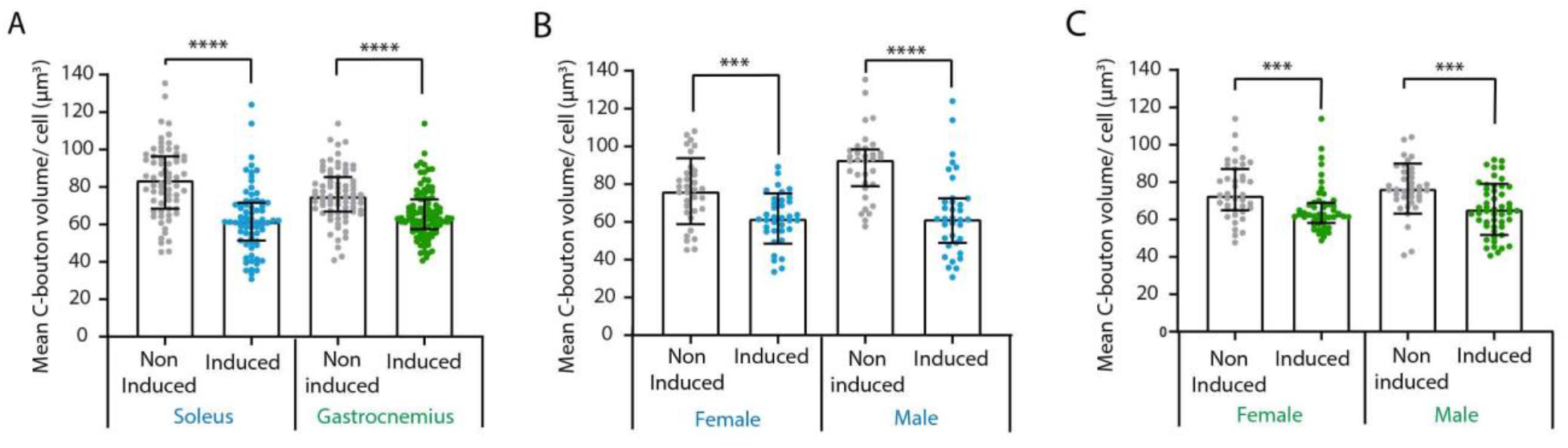
C-bouton volume is consistently reduced in induced mice. This is seen for both soleus and gastrocnemius motoneurones when the data for both sexes is collapsed (A). The decrease in C-bouton volume is consistent across the sexes for both soleus (B) and gastrocnemius (C) motoneurones.

## Discussion

The results of our study have confirmed changes in both the number and volume of C-boutons in a sporadic mouse model of ALS that aims to recapitulate the main pathological feature seen in the disease (Neumann *et al*., 2006; Scotter *et al*., 2015; Gao *et al*., 2018; Prasad *et al*., 2019). We have shown that changes in the number of C-boutons are in opposing directions on the more vulnerable, predominantly fast gastrocnemius motoneurones (a decrease) and on the less vulnerable, predominantly slow soleus motoneurones (an increase). Furthermore, these changes are sex-specific, occurring in males only. However, both sexes and motor pools showed a consistent decrease in C-bouton size. This confirms that cytoplasmic TDP-43 aggregation is sufficient to drive changes in C-boutons.

As fast motoneurones and slow motoneurones have a relatively different vulnerability in this disease, the observation that their C-boutons respond differently to the cytoplasmic TDP-43 accumulation may have implications for this vulnerability. One caveat here is that with the cell shrinkage in the gastrocnemius motoneurones, this actually resulted in an overall increase in the density of C-boutons on these cells. This emphasises that changes in C-bouton frequency, volume and soma size may be inter-related, influencing one another. This makes it important to consider all three features.

A particularly interesting finding in this study was the sex-dependent results. When analysing the collapsed data, we found a significant difference in the number of C-boutons between the non-induced control mice and the induced ALS mice for soleus motoneurones but not for gastrocnemius motoneurones. However, when splitting up the data based on sex, a significant change could now be observed for the induced male mice for both soleus and gastrocnemius motoneurones. This observation has a number of implications. First and foremost, it emphasizes the absolute necessity of not only including both sexes in such studies, but more importantly, the dangers of collapsing the data from both sexes. It suggests the possibility that some of the previous discrepancies between investigations in this area may be at least partially attributable to different proportions of each sex used. An observation of a male-specific change in C-boutons has actually already been reported in the very first mouse study in this area in SOD1 mice (Herron and Miles, 2012), yet, the vast majority of subsequent studies have failed to control for this.

Importantly, the sex difference that we have observed may have implications for sex-related differences in disease penetrance and duration. ALS occurs in men 1.5-2 times more frequently than in women (Wijesekera and Leigh, 2009; Manjaly *et al*., 2010; Brown and Al-Chalabi, 2017). While this increased risk is widely recognised in the field, we still do not fully understand the reasons behind this bias. The increased risk has resulted in a tendency for only male models of the disease to be investigated. When comparing men and women in clinical studies, men also often predominate in younger groups of the disease (McCombe and Henderson, 2010; Brown and Al-Chalabi, 2017). A sex difference is also seen in the mouse ALS models, where male mice show earlier disease onset than female mice in SOD1 models (Pfohl *et al*., 2015). The reasons behind the accelerated progression in males compared to females has yet to be determined. Oestrogen could be presumed to exert a neuroprotective effect on the motoneurones, as studies have shown that oestrogen exposure decreases the risk of developing ALS (de Jong *et al*., 2013; Rooney *et al*., 2017). Animal studies have further showed that ovariectomy in the G93A SOD1 model advances the disease progression (Yan *et al*., 2018) and previous studies have found that oestrogen exerts both a neuroprotective, anti-inflammatory and anti-apoptotic effect (Straub, 2007; Sales *et al*., 2010).

Our observed male-restricted increase in the number of C-boutons on soleus motoneurones and an increased density of C-boutons on the gastrocnemius motoneurones may further exacerbate the increased excitation of these motoneurones in males. Alternatively, it is possible that the sex differences that we have observed may simply be a result of dynamic changes in C-boutons throughout disease progression, reflecting a more advanced disease stage in the males at the same post-induction time-point. Longitudinal studies will be necessary to determine if this is the case as well as controlled survival trials with a larger cohort of mice to determine if there are sex-specific differences in the response to cytoplasmic TDP-43 deposition.

The results of this study confirm that the ectopic accumulation of cytoplasmic TDP-43 is sufficient to drive plastic changes in C-boutons. This is important as almost all of the work to date in animal models of the disease have focused on the G93A SOD1 model, which lacks the TDP-43 proteinopathy observed in most ALS patients (Suk and Rousseaux, 2020). How then, can cytoplasmic TDP-43 drive structural changes in these synapses? Normally, when located in the nucleus of healthy cells, TDP-43 regulates the expression and splicing of a wide range of proteins needed for all steps in the cell cycle. In regulating everything from mRNA transport, RNA metabolism, microRNA maturation and stress granule formation (Prasad *et al*., 2019), it shuttles between the nucleus and the cytoplasm of the cell (Pinarbasi *et al*., 2018). TDP-43 knockdown has been shown to result in significant anatomical changes and functional deficits in excitatory synapses in the hippocampus, consistent with a loss of function of the TDP-43 in driving synaptic changes (Ni *et al*., 2021). This would be consistent with the observed loss of C-boutons on gastrocnemius motoneurones, but would not explain the observed increase on soleus motoneurones. On the other hand, there is also evidence pointing towards the cytoplasmic accumulation of TDP-43 conferring a toxic gain of function (Wobst *et al*., 2017; Prasad *et al*., 2019).

In speculating how the cytoplasmic TDP-43 may drive C-bouton changes, there are two avenues to explore. The first is that, when mislocated, the TDP-43 is functionally impaired in its role in supporting adult synapses. However, this would not explain the observed increase in C-boutons. Alternatively, it is plausible that the changes in C-boutons are downstream homeostatic responses to other changes in both intrinsic and network excitability involving motoneurones or a response of the cells to the axonal problems and neuromuscular deficits seen early in the disease (Dadon-Nachum *et al*., 2011).

We believe that the earlier loss of the fast motoneurones in this disease is unlikely to explain our observed changes for a number of reasons. First, no significant cell loss has been reported in this model at 4 weeks post induction (Walker *et al*., 2015). Second, this could also not explain the increase in C-bouton number on soleus motoneurones (which are less vulnerable) or the observation that the ranges were not significantly different between our control and induced groups. Although the somas appear not to be lost at this point, it must be acknowledged that significant denervation has occurred (Walker *et al*., 2015; Spiller *et al*., 2016). Decreases in both the number and size of C-boutons are seen following axotomy and a decrease in C-bouton size has been reported after 2 weeks of impaired neuromuscular transmission (Jensen *et al*., 2020b). Although we aimed to minimize this using neuromuscular injection for the tracing, it is still possible that some neuromuscular detachment may have occurred after the tracer injection, before the animal was perfused. This may have contributed to the reduction in C-bouton size, however, it is unlikely to explain the gain of C-boutons seen on soleus motoneurones.

It is possible then that changes in either the intrinsic excitability of the spinal motoneurones or changes in the circuitry in which they are imbedded may be driving homeostatic plastic changes in the C-boutons to maintain function with disease progression. We observed that gastrocnemius motoneurone soma size decreases with disease progression, which can be predicted to alter some of the passive electrical properties of the membrane such as input resistance, making the cells easier to excite. C-boutons increase excitability by reducing the after-hyperpolarisation via reducing potassium currents through calcium dependent potassium (SK) channels and via interactions with voltage-dependent potassium (Kv2.1) channels. Therefore, an increase in C-bouton activity would be predicted to result in a further increase in excitability while a decrease would be expected to result in the converse (Miles *et al*., 2007; Romer *et al*., 2019; Nascimento *et al*., 2020). An increase in C-bouton number or density would therefore contribute to possible excitotoxicity, whereas a decrease could be interpreted as a compensatory protective mechanism, which would protect the cell from an increased excitatory drive. This of course only comes into effect if no changes are appearing at the synaptic site. To answer this question, investigations are necessary with respect to the clustering of Kv2.1-channels, SK2/3-channels and m2-receptors in this model. Future recordings from the motoneurones in the TDP-43ΔNLS animals will be necessary to test some of these hypotheses.

Recordings from cortical slice preparations from the same TDP-43ΔNLS model have demonstrated an increase in the intrinsic excitability of cortical neurones, which would increase the excitatory descending drive to the lower spinal motoneurones (Dyer *et al*., 2021). Whether the cytoplasmic TDP-43 drives similar changes in spinal motoneurones is unknown as recordings of spinal motoneurones have also largely focused on SOD1 models. Although, in ALS patients, indirect investigations using transcranial magnetic brain stimulation combined with peripheral nerve excitability tests suggest that increases in cortical excitability precede increases in spinal motoneurone excitability (Brunet *et al*., 2020). However, an increased excitability of the cortex does not necessarily translate to an increased drive to the spinal motoneurones as axonal dieback of corticospinal axons is also occurring, and so increases in the number or density of C-boutons may function to amplify a reduced drive to the lower motoneurones to maintain motor function (Brown and Al-Chalabi, 2017).

Whether our observed changes represent a homeostatic response to maintain motor output or a pathological feature contributing to potential excitotoxicity, is an open question. Attempts to genetically silence C-boutons in the G93A SOD1 mouse model have not shown any changes in survival time (time to humane endpoint). Although, the two most recent studies have reported that the mice retained better motor function (Konsolaki *et al*., 2020; Wells *et al*., 2021) and have more preserved fast twitch muscle innervation after C-bouton silencing (Wells *et al*., 2021). This suggests that targeting C-boutons may provide functional benefits to patients, although the effects may be limited. It is therefore important to determine if similar - or even more pronounced - effects can be seen in sporadic models.

Here, our results are important as a first proof of principal that C-bouton changes are not restricted to familial (SOD1) forms of the disease and that cytoplasmic TDP-43 accumulation can drive C-bouton plasticity. One caveat here is the confound of the high over-expression of TDP-43 in this model (Walker *et al*., 2015). It will therefore be important to determine if overexpression itself induces similar changes and importantly, whether an expression level, similar to that seen clinically will be sufficient to drive C-bouton changes. Here, recent advances in modelling ALS using C9orf72 repeat expansions (the most common mutations in both familial and sporadic ALS) may prove a particularly crucial development, as C9orf72 models now exist that not only develop an ALS phenotype, but also recapitulate the TDP-43 proteinopathy seen in patients (Smith *et al*., 2013; Cook *et al*., 2020). While experiments in these mice could demonstrate if C9orf72-driven levels of TDP-43 pathology is sufficient to drive C-bouton changes, for now, our results demonstrate a causal link between the TDP-43 pathology itself and the synaptic changes.

## Acknowledgments

All imaging was performed at the Core Facility for Integrated Microscopy, Faculty of Health and Medical Sciences, University of Copenhagen. ANB received a scholar stipend from the Independent Research Fund Denmark. This research was funded by The Independent Research Fund Denmark and the Læge Sofus Carl Emil Friis og Hustru Olga Doris Friis Legat.

